# Skeletal morphology of bird wings is determined by thermoregulatory demand for heat dissipation in warmer climates

**DOI:** 10.1101/2023.02.06.527306

**Authors:** Brian C. Weeks, Christina Harvey, Joseph A. Tobias, Catherine Sheard, Zhizhuo Zhou, David F. Fouhey

**Affiliations:** School for Environment and Sustainability, University of Michigan; Ann Arbor, USA; Department of Mechanical and Aerospace Engineering, University of California Davis; California, USA; Department of Life Sciences, Imperial College London; London, UK; School of Earth Sciences, University of Bristol; Bristol, UK; Department of Computer Science and Engineering, University of Michigan; Michigan, USA; School of Computer Science, Carnegie Mellon University; Pennsylvania, USA

**Keywords:** Allen’s Rule, heat dissipation, morphology, flight, wing evolution

## Abstract

The tendency for animals in warmer climates to be longer-limbed (Allen’s Rule) is widely attributed to the demands of thermoregulation. However, the underlying mechanism remains unclear, because variation in limb-length can typically be driven by selection for both efficient heat retention and increased heat dissipation capacity. Using comparative phylogenetic models, we find that occurrence in warmer climates is associated with longer wing bones for 1,520 species of passerine birds. The highly vascularized musculature along these bones is only uncovered during flight, when the wings function as the primary site of heat exchange, cooling the organism by dissipating excess heat generated by muscular activity. Conversely, the musculature along the wing bones is insulated by feathering when at rest, playing a negligible role in heat retention, even in colder climates. Given this asymmetry in thermoregulatory roles, we can identify the positive relationship between temperature and wing bone length as a phenotypic gradient shaped by increased demand for heat dissipation in warmer climates. Our findings provide a clear illustration of the mechanism by which global warming can drive spatial and temporal trends in appendage length, and also highlight the role of heat dissipation in reshaping even the most critical features of vertebrate anatomy.

**Significance Statement:** Animals tend to be longer-limbed in warmer climates, but it remains unclear whether this pattern is driven by selection for cold tolerance at low temperatures or efficient heat dissipation at high temperatures. We show that for 1,520 species of passerines, bird wing bones are relatively longer in warmer climates. The vascularized musculature along these bones primarily functions in heat exchange during flight, when the overwhelming thermoregulatory challenge is dissipating heat, suggesting longer wing-bone length is driven by heat dissipation demands. Our findings reveal the pervasive impacts of thermoregulatory demands on even the most important functional traits.

## Introduction

The need to balance energy production with heat dissipation is a fundamental imperative for endotherms. Animal species have overcome this thermoregulatory challenge in numerous ways, including via morphological adaptations to climatic niches. These adaptations form the basis for classic ecogeographic patterns, including the tendency to have smaller bodies [Bergmann’s Rule (1)] or longer limbs [Allen’s Rule (2)] in warmer climates, both of which have been reported in a wide range of animals (3–6), including humans (7). While the standard explanation for these patterns is that body size and appendage length are shaped by selection for either cold tolerance or heat dissipation, their mechanistic basis remains unclear. Unraveling these selective mechanisms is an important step towards a more complete understanding of morphological evolution (8), constraints on energy expenditure (9), and the potential utility of spatial correlations between morphology and temperature as predictors of the impacts of global warming (10).

Focusing on Allen’s Rule, we ask whether demand for shedding excess heat – as opposed to conserving heat – can drive large-scale trends in appendage length, specifically in the avian wing. Much of what we know about the effects of temperature on appendage length comes from studies of the beaks and leg bones of birds (11). There is a limit to what these appendages can tell us, however, because they regulate both heat dissipation to prevent overheating (12) and to improve cold tolerance (13), complicating the identification of underlying mechanisms. A more mechanistic understanding can be achieved by focusing on the vascularized components of wings.

In birds, the high energetic demands of flight require a 10–23-fold increase in metabolism (14), and the underlying conversion of chemical energy to mechanical energy results in nearly 90% of the metabolic energy expended during flight being converted to heat (15). Consequently, during prolonged flights, less than 1% of the heat generated by a bird is retained (16). This presents a critical thermoregulatory challenge: during flight much more heat is generated than when at rest, and nearly all of it must be dissipated to the environment. The requisite increase in heat dissipation is primarily achieved through convective heat loss from the vascularized brachial areas of the wings (17). This is in stark contrast to the negligible amounts of heat lost from the wings when individuals are not flying, and may need to retain heat, as the wings are held against the torso with the vascularized tissue well insulated under a dense layer of plumage (18).

We characterize wing heat dissipation capacity for 1,520 species of passerine birds (∼33% of all passerines) using a neural network (19) to measure key skeletal components of the wing – the humerus and the ulna – on digital photographs of 7,366 museum skeletal specimens originating from all continents except Antarctica (Fig. 1). The lengths of these wing bones determine the extent of the vascularized brachial regions of the wings, which are the main windows of heat exchange with the environment during flight (17), and are therefore expected to determine the heat dissipation capacity of the wings (Fig. 2 and Fig. S1). We couple these skeletal data with estimates of primary flight feather length and body mass (20) to quantify macroecological patterns in wing structure. Specifically, we test the hypothesis that hotter temperatures are associated with longer wing bones, driven by demand for increased heat dissipation capacity in warmer climates.

**Figure 1.**
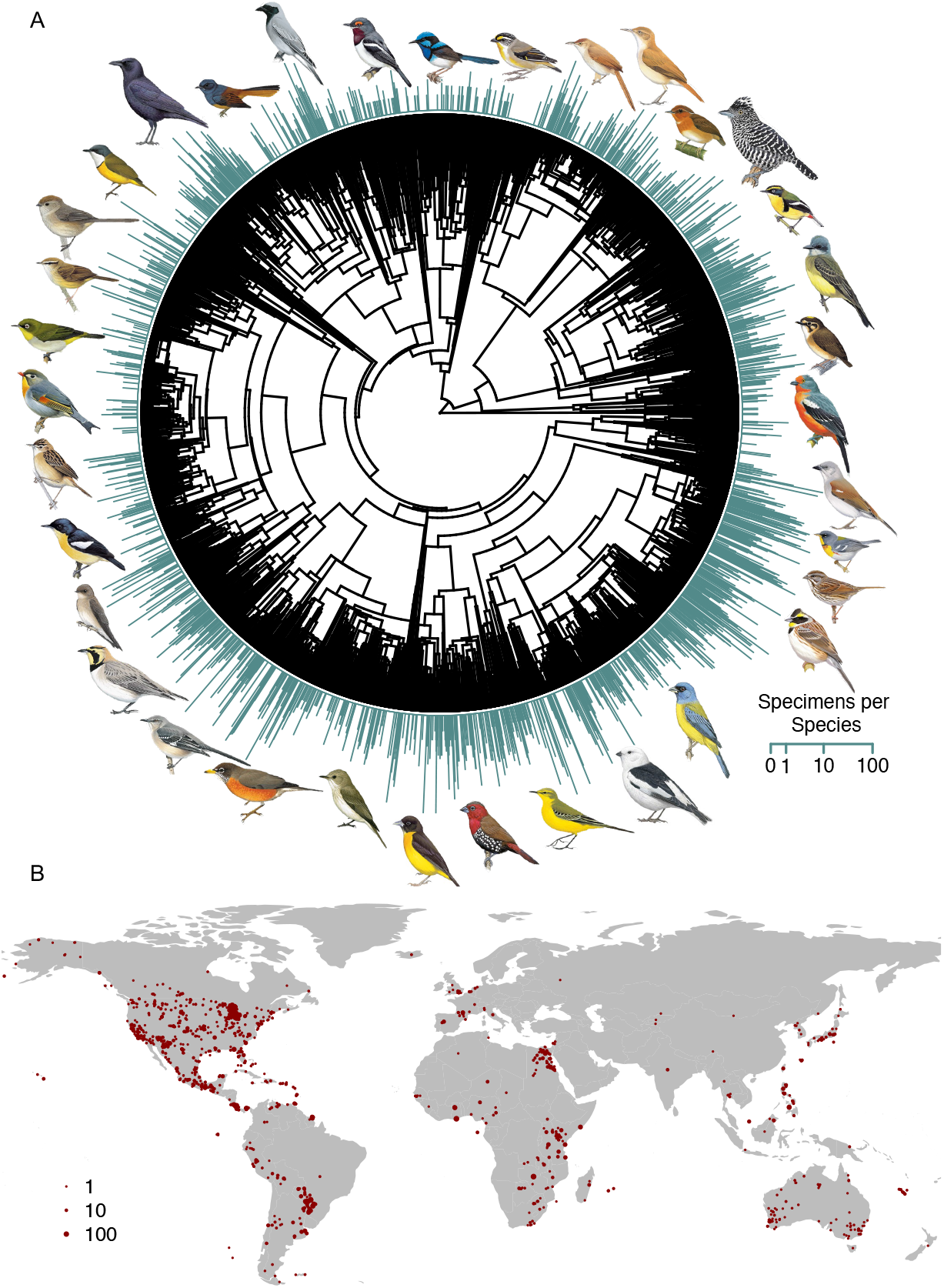
Geographic and phylogenetic sampling of avian wing-bone morphology. Using measurements from 7,366 skeletal specimens, we analyze variation in wing-bone length across 1,520 bird species, representing ∼33% of passerine diversity. (**A**) Blue bars at branch tips show the number of specimens measured per species. (**B**) Red dots show the geographic distribution of specimen localities. Images reproduced with permission from Cornell Lab of Ornithology.

**Figure 2.**
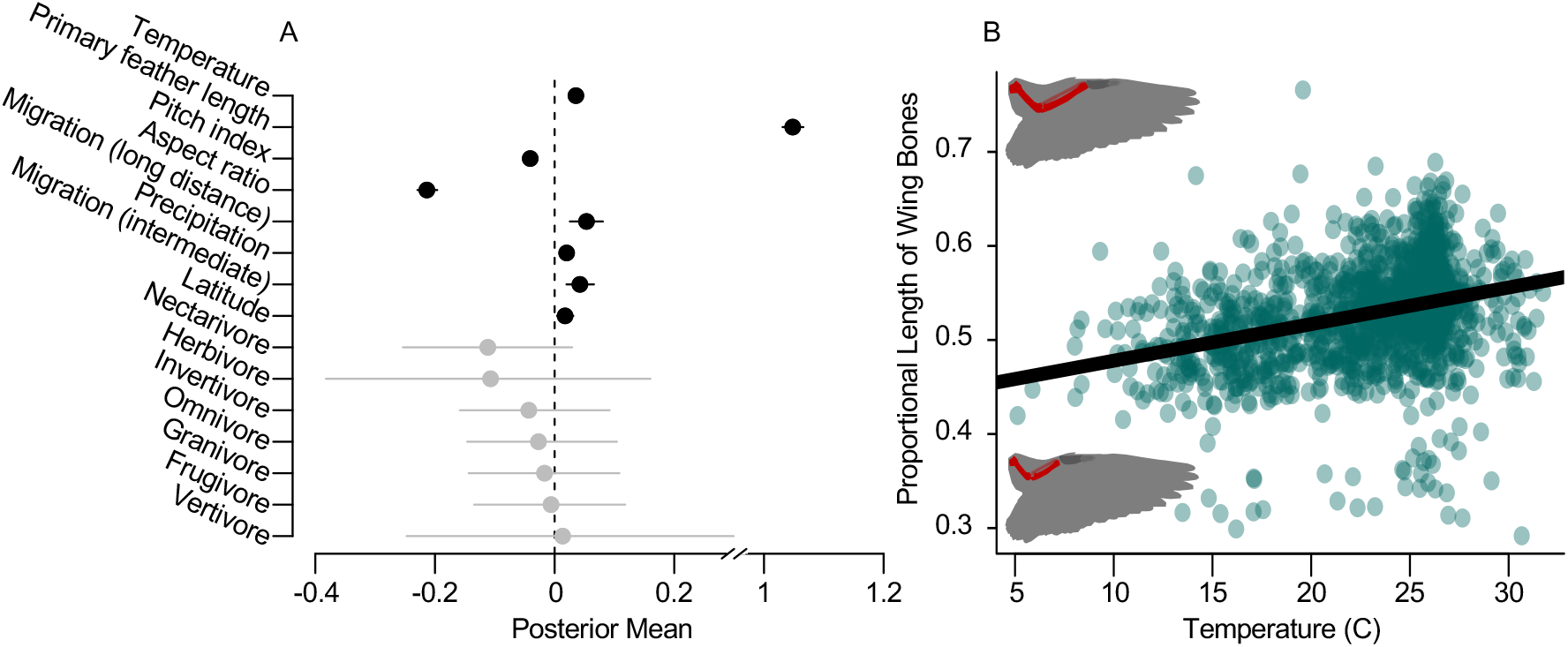
Higher temperatures are associated with increased length of avian wing-bones. (**A**) Temperature during the warmest quarter of the year is significantly positively associated with wing-bone length after controlling for the length of the primary flight feathers and a range of ecological and geographic variables (*Methods*). Dots show means of the posterior distribution with bars indicating the 95% confidence interval for each estimate; those not overlapping zero are shown in black and those overlapping zero are shown in grey. (**B**) The vascularized section of wings is relatively large in warmer climates. The proportion of the wing that is vascularized (the sum of humerus and ulna length divided by the length of the primary flight feathers) is higher for species in warmer areas. Dots represent species averages for each of the 1,502 species used in our models; a linear model of the bivariate relationship between temperature and the proportional length of the wing bones is shown.

## Results

After controlling for the range of geographic and aerodynamic variables that are expected to influence wing structure, we find a significant positive relationship between temperature and the length of the wing bones (β = 0.05; *P* < 0.001; Fig. 2). We are unlikely to have missed any alternative covariate because our model has high explanatory power, explaining nearly all variation in wing-bone length (marginal *R*^*2*^ = 0.97; conditional *R*^*2*^ > 0.99). We conclude that higher temperatures are strongly associated with longer wing bones, as predicted by Allen’s Rule.

The thermoregulation-associated pressures shaping wing structure play out within a selective landscape dominated by the demands of flight. The length of the wing bones is strongly predicted by scaling relationships, including a significant and strong positive relationship between wing-bone length and the length of the primary flight feathers (β = 1.05; *P <* 0.001; Fig. 2). The length of the primary flight feathers is highly correlated with mass (Pearson’s Correlation *(R) =* 0.88), and when mass is included as a predictor in the model rather than primary flight feather length, mass is similarly positively associated with wing bone length (β = 1.13; *P <* 0.001; Table S4). Thus, wing bones are longer in larger birds which have longer primary flight feathers.

Variation in the length of the wing bones relative to the length of the primary flight feathers is expected to impact flight dynamics. To control for the impacts of the demands of flight on wing bone length, we apply aerodynamic theory to estimate a “pitch index”, a modified index of pitch agility (21) that measures the ability of a bird to generate angular acceleration around the pitch axis (i.e., longitudinal maneuverability; Fig. S1). We also calculate the aspect ratio of the wings, a determinant of gliding flight efficiency (15, 22, 23). By controlling for pitch index and aspect ratio, we account for pressures associated with maneuverability and flight efficiency on the length of the wing bones. We find that pitch index is significantly negatively associated with wing-bone length (β = -0.04; *P <* 0.001). Wing aspect ratio is similarly significantly negatively associated with wing-bone length (β = -0.23; *P* < 0.001).

We also control for differences in migratory strategy, as long-distance movements are expected to exert strong selective pressure on wing shape and length (24). We find that intermediate- and long-distance migration, both common life history strategies in birds adapted to highly seasonal environments (25), are positively associated with the length of the wing bones (intermediate: β = 0.04; *P* < 0.001; long distance: β = 0.05; *P* < 0.001).

Finally, given the range of important ecological factors that vary with latitude, such as ecological specialization (26), species richness (27), dispersal ability (24), and geographical range size (Rapoport’s Rule), we included latitude as a predictor in our model, based on the centroid of each species’ range. We find that the distance of this centroid from the equator (absolute value of latitude) is positively associated with wing-bone length (β = 0.02; *P =* 0.01). This counter-intuitive relationship may be partially explained by migration, as our metric of migration is intended to capture major differences in life history and flight demands rather than more nuanced differences in migratory distance that may correlate with latitude and be associated with wing morphology (28). Latitude is also correlated with the quotient of wingspan divided by mass (*R* = 0.18, *P* < 0.001), and while mass is highly correlated with wing feather length (*R =* 0.88), it is not perfectly correlated. Thus, latitude may be capturing some variation in wing bone length that is driven by mass. In accordance with this idea, when mass is included in the model rather than primary flight feather length, latitude is no longer significantly associated with wing bone length (*SI*).

## Discussion

The wings of flying vertebrates are generally assumed to be shaped exclusively by the demands of aerodynamics and flight efficiency (24, 29). Our results add a new dimension by revealing a significant positive relationship between temperature and wing bone length in a broad sample of passerine birds. This pattern is consistent across a range of temperature metrics, as well as robust to statistical controls for allometry (*SI*) and a suite of ecological, geographic, and aerodynamic variables expected to influence wing structure. The anatomical trend we detect across global climatic gradients supports the hypothesis that thermoregulatory demands may be a widespread and overlooked driver of bird wing evolution.

We interpret the relationship between temperature and wing bone length as an outcome of selection for increased capacity to release heat into the environment in warmer climates because wings only become meaningful sites of heat exchange during flight (17, 18). A flying bird needs to release large amounts of body-heat to avoid over-heating and to permit the sustained energy expenditure required to maintain flight. Our findings therefore align with heat dissipation limit theory (9), and suggest that this mechanism can help to explain the widespread correlation between higher temperature and longer limb-length in animals (Allen’s Rule).

Variation in thermoregulatory demands may also explain widespread increases in the wing lengths of migrant and resident birds, despite ongoing reductions in body size over recent decades (30, 31). The opposing evolutionary trajectories of wing length and body size have prompted much debate, with the unexpected change in wing morphology mainly attributed to the need for increasingly efficient flight (30, 31). Our results provide new insight, suggesting that while rising temperatures appear to drive reduced body size in line with the expectations of Bergmann’s Rule, similar demand for efficient heat exchange also selects for increased size of wing bones, including the carpal and “hand” bones, which contribute to the measures of increased wing length reported by previous studies (30, 31).

Nearly all aspects of bird wing size and shape, including the structure of the forelimb, are expected to be broadly important to the physics of flight (22). Accordingly, we find a strong positive relationship between wing bone length and primary flight feather length (Fig. 2), reflecting the hard constraint on total wing length necessary to achieve flight that is imposed by body size (32). We also find significant negative relationships between aspect ratio and wing bone length, and between pitch index and wing bone length. This suggests that gliding flight efficiency and maneuverability are maximized in wings with relatively short wing bones compared to the primary flight feathers. These non-thermal selective pressures on wing structure likely interact via trade-offs with thermal mechanisms to determine the outcome of wing evolution.

Our results reveal that bird wings are complex traits shaped not only by flight adaptations but also by the imperative for efficient thermoregulation. The connection between wing bone length and temperature remains strongly significant even when we account for allometry, and a full range of other contextual factors, such as ecological niche (primary food source), latitude, and migratory strategy (Fig. 2). We also use aerodynamic theory to control for selection associated with flight performance (21, 22), and further account for flight performance-associated drivers of wing structure by constraining our data to passerines, thus avoiding potential confounding effects related to coarse differences in flight style on wing morphology (33). For example, heat production during flight may be significantly reduced in birds that habitually soar or glide, but this factor is removed in passerines, all of which use flapping flight. For these reasons, our models achieve unusually high explanatory power (conditional *R*^2^ > 0.99), suggesting that we are not overlooking any other major aerodynamic or ecological factor driving variation in wing-bone length.

Previous studies have shown that thermoregulatory demands can affect morphological traits functioning in food acquisition (34), with consequences for individual fitness (35). However, these studies are inconclusive regarding the relative roles of heat dissipation versus heat retention. Our analyses extend these previous findings by confirming that heat dissipation is a key mechanism driving phenotypic responses to warming temperatures, even altering the design of anatomical features fundamental to locomotion, including flight. In the case of wing morphology, the extent to which higher temperatures can lengthen wing bones is limited by biophysical constraints on organismal performance. Further analyses are needed to understand how this trade-off between thermoregulation and flight performance will impact the capacity of birds and other volant animals to respond to global warming via morphological adaptations.

## Materials and Methods

### Morphological and life history data

We estimated the length of the humerus and ulna for 7,366 individual birds of 1,520 species using skeletal specimens in the University of Michigan Museum of Zoology (UMMZ). Each specimen was photographed from a consistent distance and then measurements of the ulna and humerus were generated from the photographs with a neural network-based computer vision approach (19). In brief, for this method, training data were generated by hand annotating a subset of the images of specimens using the VGG Image Annotator (36) and the Toronto Annotation Suite (37), and then precise pixel-wise segmentations for bones of interest were annotated using AI-assisted annotation tools in the Toronto Annotation Suite (37). These partial annotations (pixel segmentations were only provided for the bones of interest) were then extended to fully annotated specimens using machine learning. Specifically, a U-Net (38) was trained to determine if each pixel was a bone or not based on preliminary annotations as well as additional synthesized samples. The U-Net was used to infer labels of all pixels in the annotated images. This annotated dataset was then used to train an instance segmentation neural network [Mask-RCNN (39)] to segment the bones. From these segmentations, the length of each bone of interest was estimated as the length of the longest diagonal. The metric size of the bone was determined using its known distance from the camera on a flat surface. For complete details, and methodological validation, see (19). We photographed all of the passerine specimens at the UMMZ, but filtered our data to only include estimates for bones that the network detected with high confidence, which we define as a detection where the most likely bone class has at least a 95% probability (19). For each specimen, if multiple measurements of the humerus or ulna were available (i.e., if the model identified and measured the bones from both wings), the arithmetic mean of the measurements was used.

Estimates of the relaxed wing chord (the distance from the carpal joint to the tip of the unflattened longest primary), which we refer to as primary flight feather length, and the distance from the carpal joint to the longest secondary flight feather were recently measured for all birds (20) and are publicly available for all of our species. Migratory status data are publicly available (20), with each species classified as 1 = non-migratory, 2 = partial/intermediate migrant, or 3 = long distance migrant. Species mean mass data and the latitude of the midpoint of the breeding range centroid were also obtained from (20).

### Environmental data

Temperature and precipitation data were generated by overlapping species breeding and resident ranges obtained from Birdlife International with a 2.5-minute resolution gridded map of the mean temperature of the warmest quarter and mean annual precipitation averaged across 1970-2000 using WorldClim v.2.1 (40), and calculating the mean values of these variables within each species’ range. All models were also refit using the 1970-2000 annual mean temperature and the maximum temperature of the warmest month (40) (SI). Species were categorized as occupying 1 of 10 ecological niches: vertivore, invertivore, scavenger, omnivore, aquatic predator, frugivore, herbivore (aquatic), herbivore (terrestrial), nectarivore or granivore, with each species assigned to a niche if the relevant food source accounts for at least 60% of the species’ diet, based on (20). We unified our taxonomy to the Birdtree taxonomy (41) using the crosswalk in (20); only species with 1:1 correspondence (i.e. that were not differentially lumped or split by the UMMZ and Birdtree taxonomy) were included.

### Flight aerodynamics

For each species, we estimated an index of maneuverability and gliding flight efficiency. Flight maneuverability was quantified using a ‘pitch index’ that is a modified version of a pitch agility metric (21), and quantifies the capacity of a bird to generate angular acceleration. To calculate the pitch index, we quantified wing area (*S*) and a static margin index (*SMI*) as:

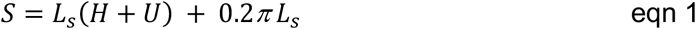

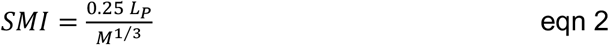

Where *L*_*s*_ is the length from the carpal joint to the tip of the first secondary feather, *H* is the length of the humerus bone, *U* is the length of the ulna bone, *L*_*p*_ is the length from the carpal joint to the tip of the longest primary feather (referred to as the relaxed wing chord) and *M* is mass. We then used these variables to calculate the Pitch Index:

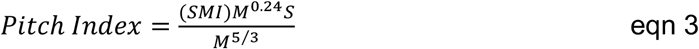

Birds with higher pitch indices have a greater ability to generate angular acceleration around the pitch axis (Figs. S1-S2). To estimate gliding flight efficiency, we calculated an index of the aspect ratio of the wings as:

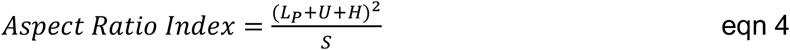

We provide derivations of these equations, empirical validation, and discussion of the necessary simplifying assumptions below.

### Statistical approach

In our models, we used species-means for all variables. Wing-bone length (the sum of the humerus length and length of the ulna) was modeled in a phylogenetic linear mixed model as a function of the length of primary feather length (i.e. the relaxed wing chord), mean temperature of the warmest quarter, the latitude at the center of the range, pitch index, aspect ratio, migration status, and ecological niche, with species identity incorporated as a random effect that accounts for phylogenetic relatedness. Mass, wing area (eqn 1), and wing loading (mass/wing area) were not included as predictors, as they were highly correlated with length of the relaxed wing chord (*R =* 0.88, 0.95, and 0.997, respectively); none of the remaining variables were highly correlated, and particularly importantly, temperature and latitude had a relatively low correlation in our data (*R = -*0.64). All continuous variables were scaled to have a mean of 0 and a standard deviation of 1 prior to model fitting; migration was included as categorical variable, with 1 being non-migratory and 3 being long distance migrants. We also constructed analogous models, controlling for log-transformed mass rather than relaxed wing chord to understand the relationship between the variables and wing-bone length relative to body size (Table S4).

Models were fit using the ‘MCMCglmm’ function in the MCMCglmm package (42) in R (43), with uninformative inverse Wishart priors (V = 1; nu = 0.02) for the residual and phylogenetic variance. To account for phylogenetic relatedness and uncertainty in those relationships, we randomly sampled 100 trees from the posterior distribution of a comprehensive phylogeny of birds (41) constructed on the Hackett et al. (44) backbone tree, and including species that were placed using taxonomy. Using a random tree from the posterior, we generated starting parameter estimates by running the models for 55,000 iterations, discarding the first 5,000 iterations as burn-in and with a thinning rate of 10. Using these starting parameter estimates, we then ran the models for 55,000 iterations, with 5,000 iterations discarded as burn-in and a thinning rate of 1,000 for each of the 100 trees. The parameter estimates were derived by combining the posterior distributions based on each of the 100 trees. To test that our results were robust to our treatment of phylogenetic uncertainty, we also fit an analogous model, but used a maximum clade credibility consensus phylogeny to capture phylogenetic relatedness and ran the model for 500,000 iterations, with the first 50,000 discarded as burn-in and a thinning factor of 100. We confirmed that the models had converged by examining parameter trace plots and posterior distributions.

## Acknowledgments

We thank Benjamin M. Winger and Brett Benz for access to skeletal specimens at the University of Michigan Museum of Zoology. This work was supported by the David and Lucille Packard Foundation.

