## Supporting Information for "Skeletal morphology of bird wings is determined by thermoregulatory demand for heat dissipation in warmer climates"

**This PDF file includes:**

Supplementary Methods

Figs. S1 to S3

Tables S1 to S5

Captions for Data S1-S3

Supplementary References

**Supplementary Methods: Derivation and Validation of the Aerodynamic Variables**

### Pitch Axis

We estimated the capacity of species to accelerate rotationally in the pitching axis with a simplified pitch index that scales with the pitch agility metric introduced by (1). Given the limited information on wing shape available for these species, it was necessary to simplify the original metric. To do this, we began by approximating the neutral point (*NP*) for each species. This approximation was informed by thin airfoil theory (2) which states that the aerodynamic center (functionally similar to the neutral point) of an airfoil is at its quarter-chord. In addition to this theoretical rationale, the neutral point has been shown empirically to scale with the quarter-chord of the standard mean chord for gull wing specimens (3). Because calculating the quarter-chord of the standard mean chord requires a complete discretization of the bird wing along its length (1), which we cannot do, we estimated the neutral point as 25% of the relaxed wing chord (i.e., the distance from the carpal joint to the longest unflattened primary feather; Fig. S2). When we regressed our estimate of the neutral point onto directly-measured data for a subset of our species, available from a previous study (1), we find that our neutral point estimate scaled with the quarter-chord of the standard mean chord (*N =* 36; *P* < 0.001; *R^2^* = 0.86).

Further, given that the location of the center of gravity along a bird’s *x*-axis cannot be differentiated from isometric scaling (mass^1/3^) (1), we estimated the ratio of the neutral point to the approximated center of gravity, hereafter known as the static margin index (SMI), as:

$SMI = \frac{NP}{M^{1/3}}$	(eqn S1)

where *M* is the full bird body mass. The higher the SMI, the more likely that the neutral point is behind the center of gravity (i.e., the bird is more stable) and vice versa. We found that the static margin index scaled with the empirically measured ratio of neutral point to the center of gravity (*N =* 36; *P* = 0.001; *R^2^* = 0.26).

Using the SMI, we generated an approximation of the pitch agility metric (1):

$Pitch Index=\frac{(SMI)M^{0.24}S}{M^{5/3}}$	(eqn S2)

where *S* is an estimate of one wing’s maximum area calculated assuming a square wing in the arm section with a width of the secondary and an elliptical wing section on the hand wing (Fig. S2). Our estimates of *S* are accurate when compared to directly-measured wing areas for a subset of our species, available from previous work (1) (*N* = 36; *P* < 0.001; *R^2^* = 0.98), and the regression relating these measures has a slope of 1.01 (Fig. S3). Note that our calculation of the pitch index assumes a cruising flight velocity estimated from previous work (4) and that the moment of inertia in the pitching axis scales isometrically (mass^5/3^) (1).

Finally, we test whether our pitch index (eqn S2) predicts previous estimates of pitch axis agility (1). We find that, as expected, pitch index scales with the capacity of a species to accelerate, quantified by the absolute maximum pitch agility metric (*N =* 36; *P* < 0.001; *R^2^* = 0.78; Fig. S3). This suggests the simplified pitch index captures similar dynamics to the pitch agility metric. In particular, birds with a higher pitch index have a larger capacity to generate a change in the pitching rate, i.e. rotational acceleration. Note that all of the above derived indices do not include the empirical constants in front of the mass terms; thus, these numbers in themselves are not directly reflective of a dimensional trait, but can be used to capture relative differences in a bird’s capacity to accelerate rotationally in the pitching axis.

The above analysis does not capture a bird’s capacity to accelerate within the roll and yaw axes. These axes are coupled both aerodynamically and inertially (5) and extracting a metric to estimate these characteristics is non-trivial. Future work is required to establish information about lateral capacity to accelerate.

### Aspect Ratio

To estimate our index of aspect ratio, we divided the square of the length of the total wing by the area of a single wing (eqn 4). We compared our estimate to direct measurements of aspect ratio for a subset of our species (6, 7), and found that our estimates are significantly correlated with published estimates (*N* = 70; *R* = 0.81; *P* < 0.001). Because our index is based on the wing area of a single wing (eqn 4), rather than total wing area, to make our estimates directly comparable aspect ratio, we divide our estimates by 2, and then regress our index onto the direct measurements of aspect ratio. We find that our aspect ratio estimates scale appropriately with aspect ratio (β = 0.86, *P* < 0.001; *R^2^* = 0.65; Fig. S3).

### Model Robustness Checks

We checked that our results were robust to different approaches for estimating temperature by fitting our model with three alternative BioClim temperature metrics: mean temperature of the warmest quarter of the year (BIO10; Fig. 3; Table S1), maximum temperature of the warmest month (BIO 5; Table S2), and mean annual temperature (BIO1; Table S3). The sign, significance, and relative strength of each parameter were consistent across models (Tables S1-S3).

In order to understand both how temperature relates to wing-bone length relative to size, and to test whether the latitude effect observed in our main model (Fig. 3) might be driven by mass, we also re-fit our main model (Fig. 3; Table S1) with the logarithm of mass as a predictor rather than relaxed wing chord (Table S4). We find that temperature remains significantly positively associated with wing-bone length (β = 0.05; *P <* 0.001), and latitude is no longer significantly associated with wing-bone length. This model is significantly less supported than the comparable model that controls for relaxed wing chord (DIC = -650 for the mass model and DIC = -1,861 for the relaxed wing chord model).

We also tested whether our treatment of phylogenetic uncertainty impacted our results by fitting the main model (Fig. 3; Table S1) but using a consensus phylogeny rather than iterating across trees from the posterior distribution of likely trees (Table S5). To do this, we built a maximum clade credibility consensus phylogeny for our species based on 1,000 posterior distributions from a comprehensive phylogeny of birds (8) constructed on the Hackett et al. (9) backbone tree, and including species that were placed using taxonomy. The consensus phylogeny was built using the ‘sumtrees’ function in dendroPy (10) following (11). The variables and priors for this model were identical to the main model (Fig. 3; Table S1), and the model was run for 500,000 iterations, with the first 50,000 iterations discarded as burn-in, and chains were thinned by a factor of 100. We find that the sign, significances, and relative strengths of all relationships are consistent when this approach is taken (Table S5).

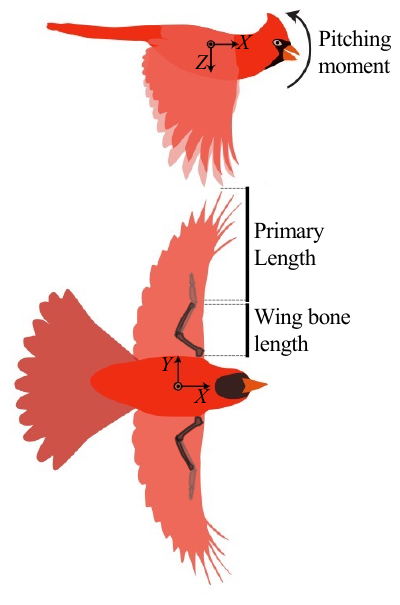

**Figure S1. Estimating maneuverability in birds.** In order to account for the demands of flight on wing structure, we control for the ability of birds to generate angular acceleration around the pitch axis (y-axis above). Roll and yaw accelerations (around the X- and Z-axes) are assumed to scale isometrically. During flight, the vascularized brachial region of the wing supported by the wing bones becomes a critical source of heat dissipation. We estimate the heat dissipation capacity of the wings using the lengths of the humerus and ulna bones (dark bones).

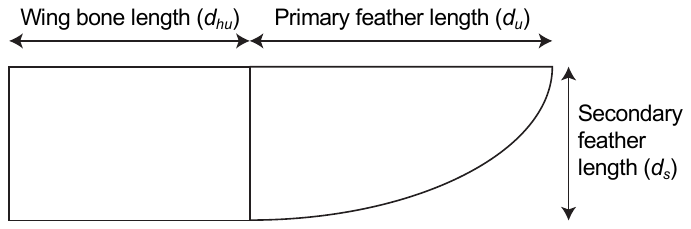

Figure S2. **Schematic diagram to illustrate method of quantifying wing structure**. To generate the pitch index, our estimate of maneuverability, we had access to the length of the wing bones (i.e. the length of the humerus plus the length of the ulna; *d_hu_*), the primary length (i.e. the relaxed wing chord, measured as the distance from the carpal joint to the tip of the longest primary feather; *d_p_*), and the distance from the carpal joint to the tip of the first secondary feather (secondary feather length; *d_s_*).

.

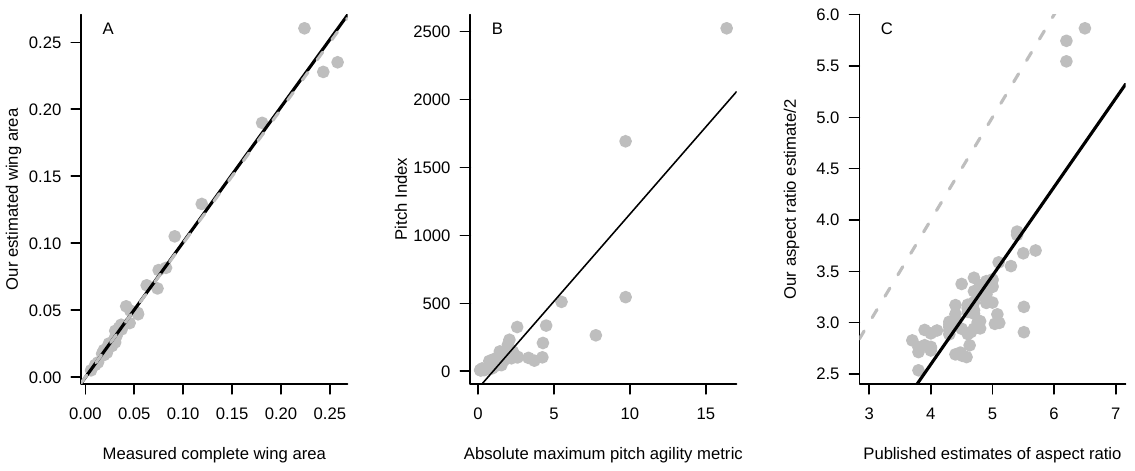

**Figure S3.** **Assessing accuracy of** wing structure and function estimates. **(A)** Wing area estimates are accurate. Comparison of wing-area estimates to directly measured wing areas from a previous study (1) shows strong clustering around the dashed gray line (1:1 relationship). A simple linear model explains most of the variation in the data (*R^2^* = 0.98; *P* < 0.001). **(B)** Our pitch index reflects pitch agility. Our pitch index estimate scales with direct estimates of the absolute maximum pitch agility metric available from (1) (linear model: *R^2^* = 0.78; *P* < 0.001). **(C)** Our aspect ratio estimates are correlated with direct measurements. For a subset of our species that had their aspect ratios directly measured in previous studies, our estimates are significantly related to the published estimates (*N =* 70; *P <* 0.001*; R^2^* = 0.65). The slope of the regression of our aspect ratio estimates (divided by 2 to make our aspect ratio index values directly comparable to aspect ratio) onto published estimates (black line) is 0.86. This is close to the 1:1 expectation (dashed gray line); the difference in intercept is not expected to influence our results as we use relative aspect ratios.

**Supplementary Tables**

In all tables, the length of the wing bones is the dependent variable, and parameters with significant relationships to the length of the wing bones (pMCMC < 0.05) are shown in bold.

Table S1. Model results predicting wing-bone length using mean temperature of warmest quarter. These results are presented in the main text (Fig. 3), with the methods and variables described in *Materials and Methods*. The DIC for this model = -1,938; G-structure (phylogeny) has a posterior mean of 0.02, with effective sample size = 337; R-structure (variable effects) has a posterior mean of 0.01, with effective sample size = 379.

| Parameter | Posterior mean | 95% CI | Effective sample size | pMCMC |
| --- | --- | --- | --- | --- |
| **Temperature of the warmest quarter** | **0.04** | **0.03 – 0.05** | **3,465** | **< 0.001** |
| **Latitude** | **0.02** | **0.004 – 0.03** | **4,700** | **0.01** |
| **Relaxed wing chord** | **1.05** | **1.03 – 1.07** | **914** | **<0.001** |
| Frugivore | -0.01 | -0.13 – 0.12 | 5,000 | 0.92 |
| Granivore | -0.02 | -0.14 – 0.11 | 5,000 | 0.79 |
| Terrestrial herbivore | -0.11 | -0.38 – 0.16 | 4,435 | 0.44 |
| Invertivore | -0.04 | -0.16 – 0.09 | 5,000 | 0.48 |
| Nectarivore | -0.11 | -0.25 – 0.03 | 4,775 | 0.13 |
| Omnivore | -0.03 | -0.15 – 0.10 | 5,000 | 0.66 |
| Vertivore | 0.01 | -0.25 – 0.30 | 5,000 | 0.94 |
| **Pitch index** | **-0.04** | **-0.05 – 0.03** | **2,160** | **<0.001** |
| **Aspect ratio** | **-0.21** | **-0.23 - -0.20** | **3,295** | **<0.001** |
| **Migration (I)** | **0.04** | **0.02 – 0.07** | **5,000** | **<0.001** |
| **Migration (LD)** | **0.05** | **0.03 – 0.08** | **4,126** | **<0.001** |
| **Precipitation** | **0.02** | **0.01 – 0.03** | **5,000** | **<0.001** |

Table S2. Model results predicting wing-bone length using maximum temperature of the warmest quarter. This model is identical to the main model (Fig. 3; table S1) except that it uses the maximum temperature of the warmest month rather than the mean temperature of the warmest quarter as a predictor. The DIC for this model = -1,936; G-structure (phylogeny) has a posterior mean of 0.03, with effective sample size = 265; R-structure (variable effects) has a posterior mean of 0.01, with effective sample size = 295.

| Parameter | Posterior mean | 95% CI | Effective sample size | pMCMC |
| --- | --- | --- | --- | --- |
| **Maximum temperature of warmest month** | **0.03** | **0.02 – 0.04** | **4,083** | **<0.001** |
| **Latitude** | **0.02** | **0.003 – 0.03** | **4,009** | **0.01** |
| **Relaxed wing chord** | **1.05** | **1.03 – 10.6** | **601** | **<0.001** |
| Frugivore | -0.002 | -0.12 – 0.13 | 5,000 | 0.96 |
| Granivore | -0.01 | -0.14 – 0.11 | 5,000 | 0.84 |
| Terrestrial herbivore | -0.10 | -0.39 – 0.17 | 5,000 | 0.45 |
| Invertivore | -0.04 | -0.17 – 0.08 | 3,799 | 0.53 |
| Nectarivore | -0.11 | -0.25 – 0.04 | 5,000 | 0.13 |
| Omnivore | -0.02 | -0.15 – 0.10 | 5,000 | 0.71 |
| Vertivore | 0.02 | -0.26 – 0.29 | 5,000 | 0.85 |
| **Pitch index** | **-0.04** | **-0.05 - -0.03** | **2,389** | **<0.001** |
| **Aspect ratio** | **-0.21** | **-0.23 - -0.20** | **3,055** | **<0.001** |
| **Migration (I)** | **0.04** | **0.02 – 0.06** | **4,183** | **<0.001** |
| **Migration (LD)** | **0.05** | **0.01 – 0.08** | **2,712** | **<0.001** |
| **Precipitation** | **0.03** | **0.02 – 0.04** | **4,559** | **<0.001** |

Table S3. Model results predicting wing-bone length using mean annual temperature. This model is identical to the main model (Fig. 3; table S1) except that it uses mean annual temperature rather than mean temperature of the warmest quarter as a predictor. The DIC for this model = -1,911; G-structure (phylogeny) has a posterior mean of 0.03, with effective sample size = 318; R-structure (variable effects) has a posterior mean of 0.01, with effective sample size = 362.

| Parameter | Posterior mean | 95% CI | Effective Sample Size | pMCMC |
| --- | --- | --- | --- | --- |
| **Mean annual temperature** | **0.04** | **0.03 – 0.06** | **4,683** | **<0.001** |
| **Latitude** | **0.03** | **0.01 – 0.04** | **5,000** | **0.002** |
| **Relaxed wing chord** | **1.05** | **1.03 – 1.07** | **854** | **<0.001** |
| Frugivore | 0.003 | -0.13 – 0.14 | 5,000 | 0.97 |
| Granivore | -0.01 | -0.14 – 0.13 | 5,000 | 0.90 |
| Terrestrial herbivore | -0.11 | -0.40 – 0.17 | 4,775 | 0.41 |
| Invertivore | -0.03 | -0.17 – 0.09 | 5,000 | 0.60 |
| Nectarivore | -0.11 | -0.26 – 0.03 | 5,000 | 0.14 |
| Omnivore | -0.02 | -0.15 – 0.11 | 5,000 | 0.77 |
| Vertivore | 0.03 | -0.26 – 0.30 | 5,000 | 0.84 |
| **Pitch index** | **-0.04** | **-0.05 – -0.03** | **3,316** | **<0.001** |
| **Aspect ratio** | **-0.21** | **-0.23 – -0.20** | **3,049** | **<0.001** |
| **Migration (I)** | **0.05** | **0.02 – 0.07** | **4,079** | **<0.001** |
| **Migration (LD)** | **0.06** | **0.04 – 0.09** | **5,000** | **<0.001** |
| **Precipitation** | **0.02** | **0.01 – 0.03** | **5,000** | **<0.001** |

Table S4. Model results controlling for body mass. This model is identical to the main model (Fig. 3; table S1) except that it uses body mass as predictor rather than relaxed wing chord. The DIC for this model = -746 (significantly higher than the model including wing length: DIC = -1,938); G-structure (phylogeny) has a posterior mean of 0.11, with effective sample size = 111; R-structure (variable effects) has a posterior mean of 0.02, with effective sample size = 83.

| Parameter | Posterior mean | 95% CI | Effective sample size | pMCMC |
| --- | --- | --- | --- | --- |
| **Mean temperature of the warmest quarter** | **0.04** | **0.03 – 0.06** | **685** | **<0.001** |
| Latitude | 0.003 | -0.02 – 0.03 | 1,101 | 0.77 |
| Mass | **1.13** | **1.09 – 1.17** | **172** | **<0.001** |
| Frugivore | -0.06 | -0.26 – 0.16 | 3,993 | 0.60 |
| Granivore | -0.03 | -0.23 – 0.19 | 4,205 | 0.80 |
| Terrestrial herbivore | -0.44 | -0.94 – 0.03 | 4,787 | 0.08 |
| Invertivore | -0.10 | -0.30 – 0.11 | 4,172 | 0.34 |
| Nectarivore | -0.09 | -0.32 – 0.14 | 4,517 | 0.45 |
| Omnivore | -0.08 | -0.29 – 0.12 | 4,071 | 0.42 |
| Vertivore | -0.18 | -0.59 – 0.26 | 5,000 | 0.40 |
| **Pitch index** | **0.35** | **0.32 – 0.38** | **430** | **<0.001** |
| Aspect ratio | 0.01 | -0.02 – 0.04 | 317 | 0.68 |
| Migration (I) | -0.005 | -0.04 – 0.03 | 632 | 0.82 |
| Migration (LD) | 0.04 | -0.001 – 0.09 | 2,955 | 0.06 |
| Precipitation | 0.003 | -0.01 – 0.02 | 1,699 | 0.73 |

Table S5. Model results using a consensus phylogeny. When accounting for phylogenetic relatedness in our model using a consensus phylogeny, rather than iterating across trees from the posterior distribution, we find that the same variables are significant and the direction and relative magnitude of the relationships are qualitatively the same.

| Parameter | Posterior mean | 95% CI | Effective sample size | pMCMC |
| --- | --- | --- | --- | --- |
| **Mean temperature of the warmest quarter** | **0.04** | **0.03 – 0.05** | **44,383** | **<0.001** |
| **Latitude** | **0.02** | **0.004 – 0.03** | **42,262** | **0.01** |
| **Relaxed wing chord** | **1.05** | **1.03 – 1.06** | **36,927** | **<0.001** |
| Frugivore | -0.01 | -0.14 – 0.12 | 43,501 | 0.87 |
| Granivore | -0.02 | -0.15 – 0.11 | 43,211 | 0.75 |
| Terrestrial herbivore | -0.13 | -0.41 – 0.15 | 45,000 | 0.34 |
| Invertivore | -0.05 | -0.17 – 0.07 | 43,211 | 0.46 |
| Nectarivore | -0.12 | -0.26 – 0.02 | 43,685 | 0.11 |
| Omnivore | -0.03 | -0.15 – 0.09 | 43,381 | 0.65 |
| Vertivore | 0.01 | -0.26 – 0.28 | 45,000 | 0.94 |
| **Pitch index** | **-0.04** | **-0.05 - -0.03** | **42,050** | **<0.001** |
| **Aspect ratio** | **-0.21** | **-0.23 - -0.20** | **35,437** | **<0.001** |
| **Migration (I)** | **0.04** | **0.02 – 0.07** | **45,000** | **<0.001** |
| **Migration (LD)** | **0.05** | **0.03 – 0.08** | **45,000** | **<0.001** |
| **Precipitation** | **0.02** | **0.01 – 0.03** | **44,263** | **<0.001** |
